# Glutamic acid is a carrier for hydrazine during the biosyntheses of fosfazinomycin and kinamycin

**DOI:** 10.1101/365031

**Authors:** Kwo-Kwang Abraham Wang, Tai L. Ng, Peng Wang, Zedu Huang, Emily P. Balskus, Wilfred A. van der Donk

## Abstract

Fosfazinomycin and kinamycin are natural products that contain nitrogen-nitrogen (N-N) bonds but that are otherwise structurally unrelated. Despite their considerable structural differences, their biosynthetic gene clusters share a set of genes predicted to facilitate N-N bond formation. In this study, we show that for both compounds, one of the nitrogen atoms in the N-N bond originates from nitrous acid. Furthermore, we show that for both compounds, an acetylhydrazine biosynthetic synthon is generated first and then funneled via a glutamyl carrier into the respective biosynthetic pathways. Therefore, unlike other pathways to NN bond-containing natural products wherein the N-N bond is formed directly on a biosynthetic intermediate, during the biosyntheses of fosfazinomycin, kinamycin, and related compounds, the N-N bond is made in an independent pathway that forms a branch of a convergent route to structurally complex natural products.

---

More than 200 natural products containing nitrogen-nitrogen (N-N) bonds have been identified with various bioactivities **(Fig. 1a)**^1^. For instance, streptozotocin, a nitrosamine-containing compound, exhibits cytotoxic activity and is currently deployed clinically^2^, and valanimycin, featuring an azoxy group, displays both antimicrobial and cytotoxic activities^1^. Other examples are azamerone containing a pyridazine, kutzneride with a piperazic acid^2, 3^, the fosfazinomycins having a phosphonohydrazide^4^, and a group of molecules including cremeomycin, the lomaiviticins and the kinamycins containing diazo groups^1^.

**Figure 1.**
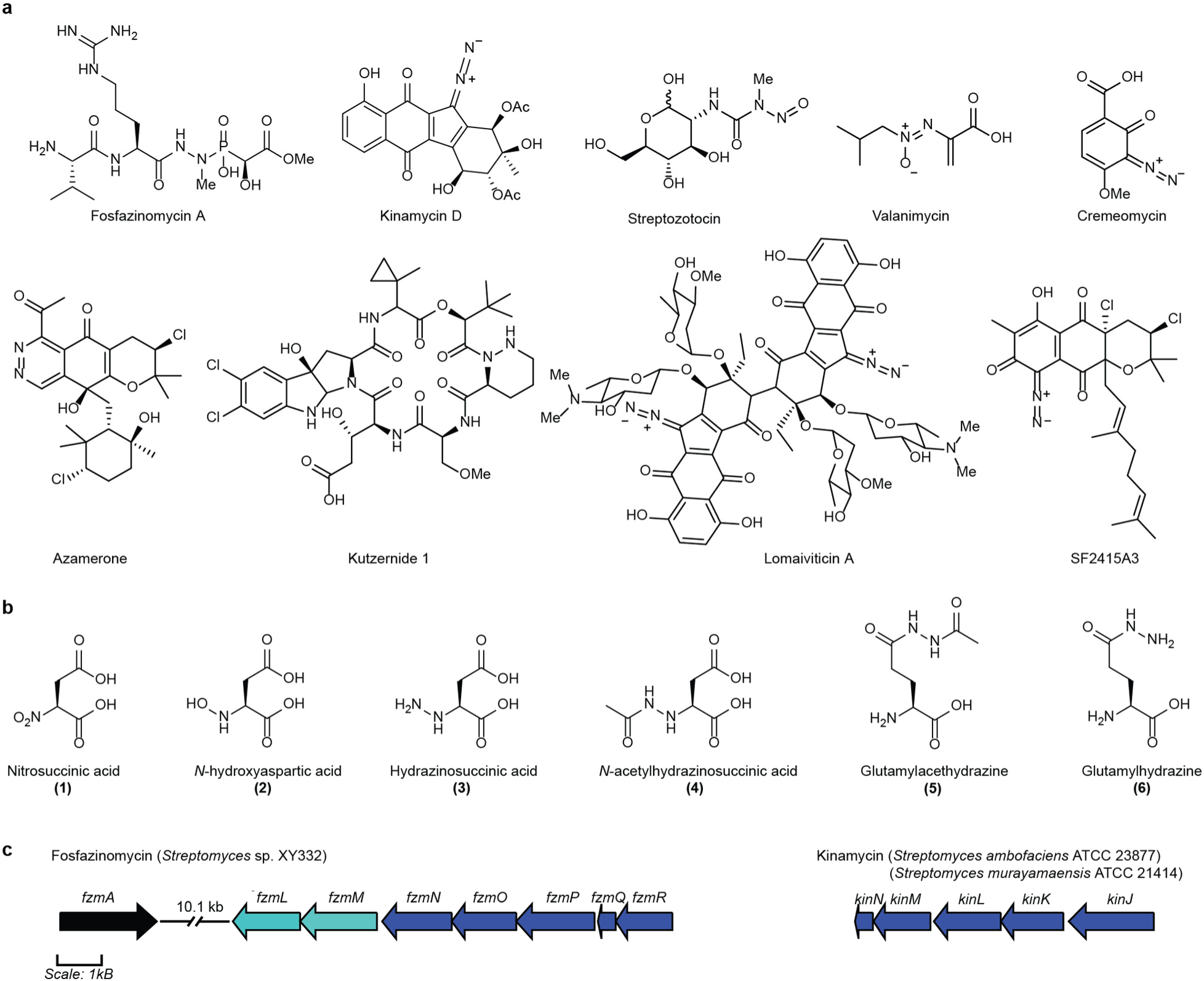
Structures and biosynthetic gene clusters of fosfazinomycin and kinamycin. **(a)** The structures of a select group of N-N bond containing natural products. **(b)** The structures of the biosynthetic intermediates discussed in this study. **(c)** Selected segments of the biosynthetic gene clusters of fosfazinomycin and kinamycin that encompass the genes discussed in this study. The five conserved genes in the two clusters are colored in dark blue. *fzmL* and *fzmM*, which encode enzymes responsible for forming nitrous acid from aspartic acid, are colored in light blue.

Despite the diversity of N-N bonds found in natural products, relatively little is currently known about their biosyntheses. Based on the available biosynthetic gene clusters, the biosynthetic logic of N-N bond formation appears to be at least partially shared among certain groups of compounds. Recently, N-N bond formation was reconstituted *in vitro* for kutzneride, showing that N5 on ornithine is oxidized to give N5-hydroxyornithine by a flavin-dependent enzyme before the N-N bond is formed by a heme protein (**Supplementary Fig. 1a**)^5^. Thus, for this compound, one nitrogen is activated to a more electrophilic species and then reacted intramolecularly with a nucleophilic nitrogen.

A different strategy towards N-N bond formation features the formal reaction of a nitrogen-containing intermediate with nitrous acid. For instance, during the biosynthesis of cremeomycin, a flavin-dependent monooxygenase, CreE, oxidizes aspartic acid to nitrosuccinic acid (**1**, **Fig. 1b**), and a lyase, CreD, subsequently liberates nitrous acid^6^. Nitrous acid is then proposed to partake in diazotization with a primary aromatic amine on an advanced intermediate to form the N-N bond, but the details of this process and whether it requires an enzyme remain unclear (**Supplementary Fig. 1b**)^7^. Recently, we reported that the fosfazinomycin biosynthetic enzymes FzmM and FzmL, homologs of CreE and CreD respectively, can also produce nitrous acid from aspartic acid with *N-*hydroxyaspartic acid (**2**, **Fig. 1b**) as an intermediate^8^. Moreover, we showed that aspartic acid is incorporated into at least one of the nitrogen atoms in the phosphonohydrazide moiety of fosfazinomycin. Interestingly, a set of five genes is conserved between the biosynthetic gene clusters of fosfazinomycin (*fzm*) and kinamycin (*kin*) (**Fig. 1c**), suggesting that the two compounds may share biosynthetic steps. These genes are predicted to encode homologs of a glutamine synthetase (FzmN/KinL), an amidase (FzmO/KinK), a hypothetical protein (FzmP/KinJ), an *N-*acetyltransferase (FzmQ/KinN), and an adenylosuccinate lyase (FzmR/KinM)^9, 10^. The roles of these five enzymes in installing the N-N bond into two structurally divergent compounds are not known. In our prior work concerning the biosynthesis of fosfazinomycin, we reconstituted *in vitro* the activities of FzmQ and FzmR. FzmQ performs the acetylation of hydrazinosuccinic acid (**3**) to yield *N*-acetylhydrazinosuccinic acid (**4**, **Fig. 1b**), and FzmR catalyzes an elimination reaction of **4** to afford fumaric acid and acetylhydrazine^8^.

In this study we show via isotope labeling experiments that one of the nitrogen atoms in the N-N bonds of both fosfazinomycin and kinamycin originates from nitrous acid. Additional feeding experiments and reconstitution of enzymatic activities confirm the intermediacies of **2, 4**, and acetylhydrazine in both biosynthetic pathways. Moreover, we demonstrate that after the N-N bond is formed, the hydrazine moiety is installed onto the side chain carboxyl group of glutamic acid. These results reveal that during the biosyntheses of both fosfazinomycin and kinamycin, the N-N bond is formed from aspartic acid as a discrete and separate synthon before being shuttled into the two respective biosynthetic pathways using glutamic acid as carrier. Thus, these pathways are fundamentally different from the biosynthetic routes to kutzneride and cremeomycin in which the N-N bond is formed directly on a biosynthetic intermediate.

## Results

### Confirmation of fosfazinomycin biosynthetic intermediates by feeding experiments

In previous work, we showed using stable isotope labeling experiments that at least one of the nitrogen atoms in the phosphonohydrazide linkage in fosfazinomycin is derived from aspartic acid^8^. Furthermore, reconstitution of the activities of FzmM and FzmL demonstrated that they produce nitrous acid from aspartic acid; however, we also found that under certain conditions, FzmM converted aspartic acid to *N*-hydroxyaspartic acid (**2**)^*8*^. In light of the recent work on kutzneride and cremeomycin, it was unclear whether the generation of **2** by FzmM is a direct intermediate in N-N bond formation (similar to kutzneride, **Supplementary Fig. S1a**) or if **2** is only an intermediate in the oxidation of aspartic acid to **1** (similar to cremeomycin, **Supplementary Fig. S1b**)^5, 7^. In order to investigate these possibilities, we performed isotope labeling studies in the native producing organism of fosfazinomycin, *Streptomyces* sp. NRRL S-149. First, the organism was cultivated in a minimal medium with ^15^NH_4_Cl and ^15^N-aspartic acid as the only two sources of nitrogen, and analysis by liquid chromatography-high resolution mass spectrometry (LC-HRMS) of the spent medium revealed a singly charged peak ([M+H]^+^ = 461.1957 Da) corresponding to fosfazinomycin A containing seven ^15^N atoms (**Fig. 2a**). Cultures grown in uniformly ^15^N-labeled medium were then fed sodium nitrite at natural isotopic abundance. LC-HRMS analysis of the spent medium revealed a clear increase in the peak corresponding to [M+H]^+^ =460.1996 Da (**Fig. 2b**), one mass unit lower than uniformly ^15^N-labeled fosfazinomycin. Integration of the peak areas showed that approximately 30% of the product contains a single ^14^N atom originating from nitrite. We next fed **2** at natural abundance to *Streptomyces* sp. NRRL S-149 grown in medium containing ^15^NH_4_Cl and ^15^N-aspartic acid as the only two other sources of nitrogen. As with nitrite, a significant peak with [M+H]^+^ = 460.1993 Da was observed (**Fig. 2c**), corresponding to a single incorporation (20%) of ^14^N into fosfazinomycin.

**Figure 2.**
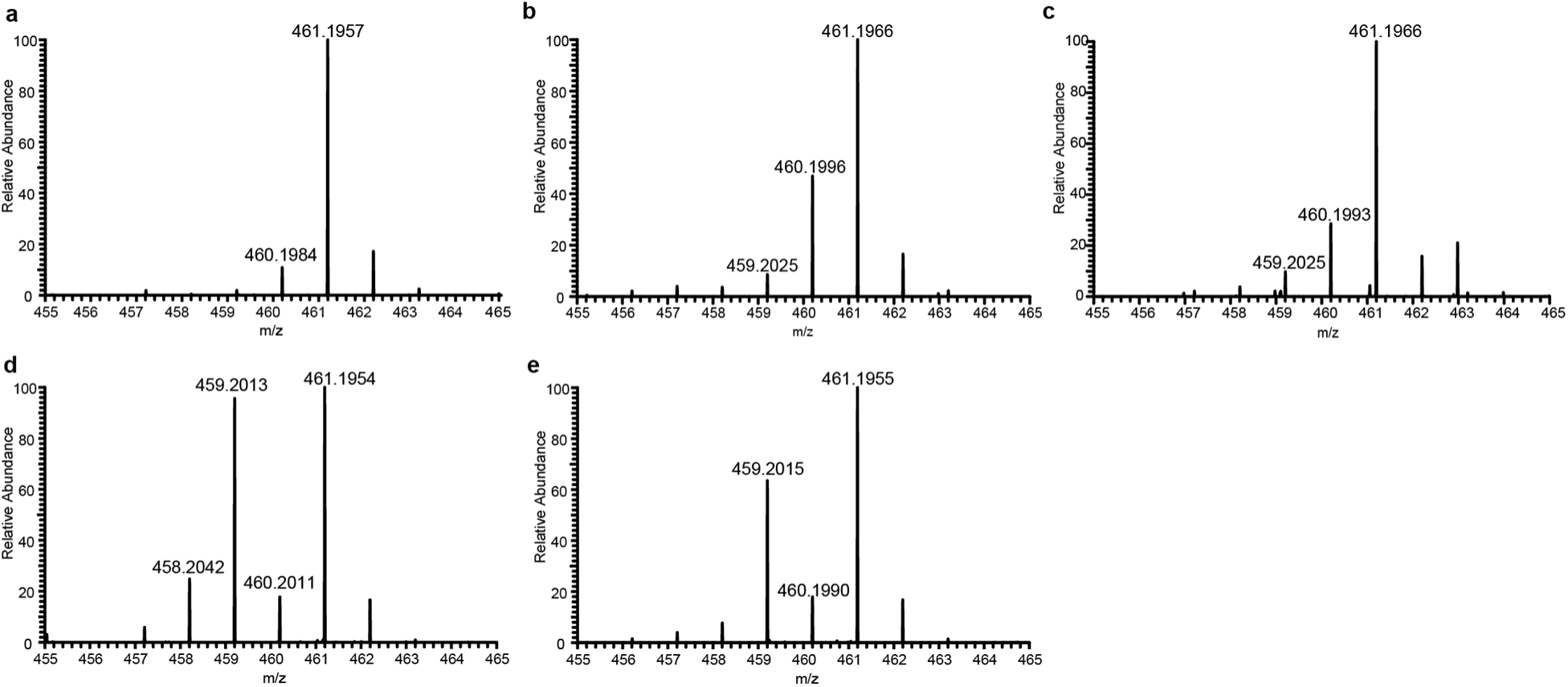
HRMS analysis of ^14^N-incorporation into ^15^N-labeled fosfazinomycin A in *Streptomyces* sp. NRRL S-149. **(a)** Fosfazinomycin A produced in uniformly ^15^N-labeled media. The calculated m/z ([M+H]^+^) for uniformly ^15^N-labeled fosfazinomycin A is 461.1966. **(b, c)** Spectra from cells cultivated in uniformly ^15^N-labeled media and fed with either **(b)** NaNO_2_ or **(c)** *N*-hydroxyaspartic acid (**2**). The calculated m/z ([M+H]^+^) for fosfazinomycin A with a single ^14^N-incorporation is 460.1996. **(d, e)** Spectra from cells grown in uniformly ^15^N-labeled media and fed with either **(d)** hydrazinosuccinic acid (**3**) or **(e)** acetylhydrazine. The calculated m/z ([M+H]^+^) for fosfazinomycin A with two ^14^Nincoporations is 459.2025.

We next performed tandem mass spectrometry (MS/MS) analysis to determine if the ^14^N atoms from nitrite and **2** were incorporated into the hydrazide linkage of fosfazinomycin. First, as a control, uniformly ^15^N-labled fosfazinomycin ([M+H]^+^ = 461.20) was subjected to higher energy collisional dissociation (HCD), and the product and fragment ions were assigned (**Supplementary Fig. 2a**). Then, the spent medum from the culture cultivated with added unlabeled sodium nitrite was analyzed focusing on a precursor ion ([M+H]^+^ = 460.20) corresponding to singly ^14^N-labeled fosfazinomycin (**Supplementary Fig. 2b**). Analysis of the fragment ions indicated ^14^N-incorporation only if the fragment contained the N-N bond (ions g, j, and k, accented by the blue stars). Likewise, in the sample fed with **2**, the singly ^14^N-labeled fosfazinomycin (precursor ion [M+H]^+^ = 460.20) was fragmented (**Supplementary Fig. 2c**). Again, only fragment ions containing the N-N bond (ions g, j, and k) incorporated ^14^N. Thus, these feeding experiments demonstrate that one of the nitrogens in the phosphonohydrazide linkage in fosfazinomycin is derived from nitrite. These data are also consistent with our previous study that showed that nitrite is formed from Asp with **2** as an intermediate.

We next performed additional feeding experiments to confirm that **3** and acetylhydrazine are also competent intermediates in the fosfazinomycin biosynthetic pathway. When **3** at natural isotopic abundance was fed to the producing organism grown in ^15^N-labeled medium, the product ions showed a large increase in a peak two mass units lower [M+H]^+^ = 459.2013) than the uniformly ^15^N-labeled product (**Fig. 2d**). The data demonstrate about 50% double incorporation of ^14^N into fosfazinomycin. Similarly, when acetylhydrazine (nitrogen at natural isotopic abundance) was added to the ^15^N-labeled medium, a large increase in a peak two mass units lower ([M+H]^+^ = 459.2015) relative to the control was observed, indicating two incorporations of ^14^N (40%) into fosfazinomycin A (**Fig. 2e**). MS/MS was then used to determine the location of ^14^N incorporation. Samples obtained from feeding **3** were analyzed by MS/MS using doubly ^14^N-labeled fosfazinomycin as the precursor ion ([M+H]^+^ = 459.20) (**Supplementary Fig. 2d**). After HCD, only the fragment ions that retained the N-N bond (ions f, g, j, and k indicated by the blue stars) exhibited a −2 Da mass shift compared to the control indicating selective incorporation of ^14^N into the N-N bond of fosfazinomycin. Likewise, when doubly ^14^N-labeled fosfazinomycin (precursor [M+H]^+^ = 459.20) obtained from feeding acetylhydrazine was fragmented, only product or fragment ions containing the N-N bond incorporated two ^14^N atoms (**Supplementary Fig. 2e**). Thus, these data strongly suggest that **3** and acetylhydrazine are intermediates in the pathway and that both of their nitrogen atoms are incorporated selectively into the phosphonohydrazide linkage of fosfazinomycin, consistent with the demonstration that *in vitro* FzmQR convert **3** to acetylhydrazine^8^.

### Feeding studies in a kinamycin producer

As described in the introduction, the kinamycin biosynthetic gene cluster contains genes that are homologous to *fzmQR*, suggesting that **3** and acetylhydrazine might also be intermediates in its biosynthesis. Furthermore, kinamycin D has been recently reconstituted *in vivo* in *S. albus* J1074, and its heterologous expression has been demonstrated to be dependent on *alp2F* and *alp2G*, orthologs of *fzmM and fzmL*^11^.Thus, to investigate whether the products of FzmLM are incorporated into kinamycin, we performed a series of stable isotope feeding studies.

We began by supplementing the fermentation medium of *S. murayamaensis*, a prolific producer of kinamycin D^12^, with 1 mM of ^15^N-nitrite (**Fig. 3a**). LC-HRMS analysis of the organic extracts from the ^15^N-nitrite-fed cultures revealed ~90% enrichment of singly labeled kinamycin D ([M+H]^+^ = 456.1055) (**Fig. 3b**). We isolated the natural product and observed by 1D ^15^N-NMR spectroscopy that the proximal nitrogen of the diazo group is exclusively enriched (**Fig. 3c**). This outcome contrasts sharply with the results of previous ^15^N-nitrite feeding experiments performed with the producers of the diazo-containing secondary metabolite SF2415A3, in which only the distal diazo nitrogen atom was labeled (**Supplementary Fig 1c)**^3^. In this study, both ^15^N-nitrite and ^15^N-nitrate selectively enriched the distal diazo nitrogen atom. We tested whether ^15^Nnitrate also labeled the diazo group of kinamycin D. After culturing *S. murayamaensis* in the presence of 1 mM ^15^N-nitrate, we also observed ~80% enrichment of the proximal nitrogen (**Fig. 3ac**). Diazo formation in the biosynthesis of kinamycin had been proposed to occur via a late-stage N-N bond formation involving an aminobenzo[*b*]fluorene biosynthetic precursor such as stealthin C, a biosynthetic logic that would parallel that of SF2415A3 (**Supplementary Fig. 1cd)**^13^. The localization of ^15^N label to the proximal diazo nitrogen atom in our feeding experiments shows that this is not the case, and supports the involvement of an alternative strategy for diazo installation in kinamycin biosynthesis.

**Figure 3.**
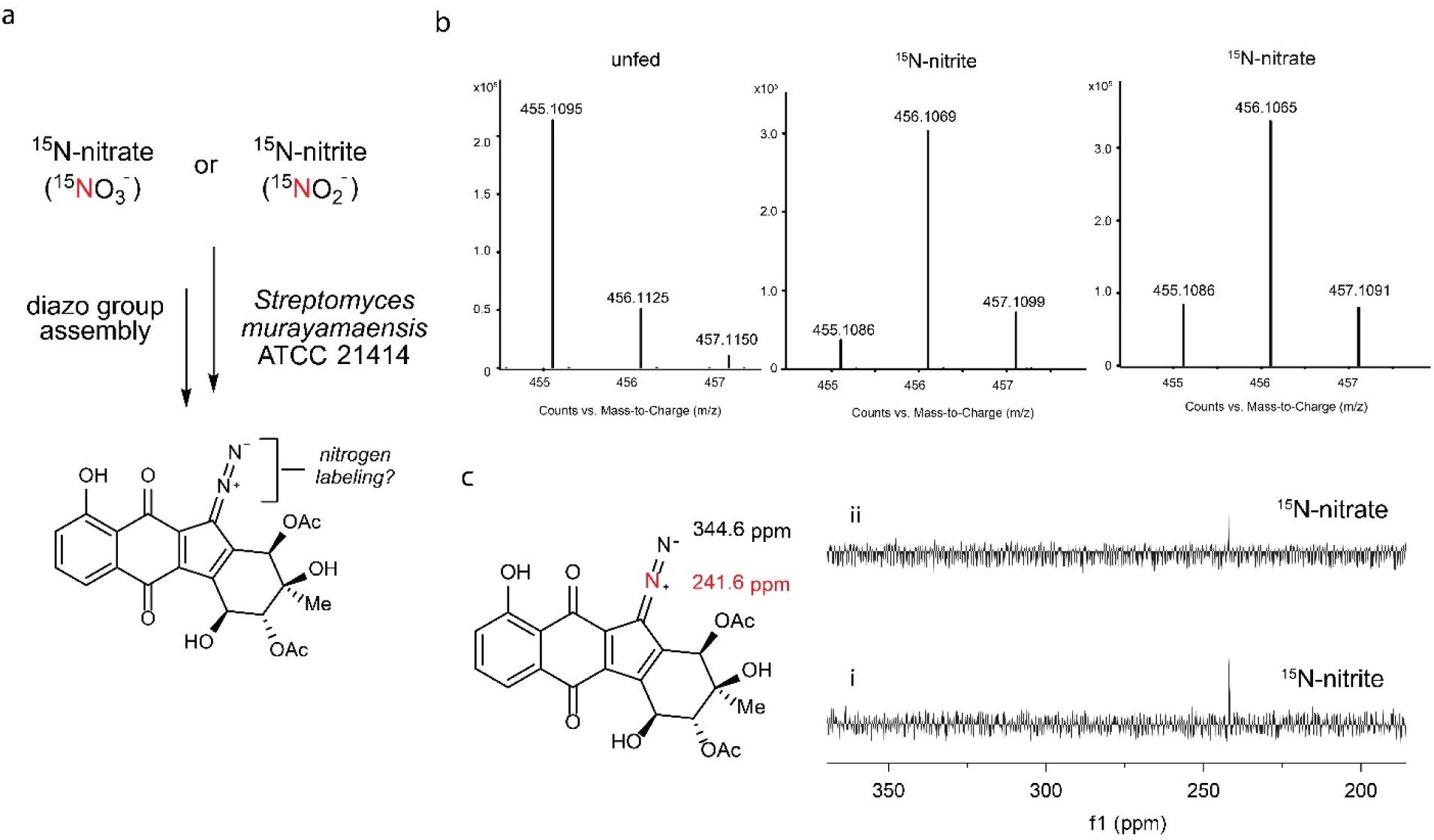
^15^N-incorporation into kinamycin D by inorganic nitrogen sources. **(a)** Scheme for feeding experiments with *S. murayamaensis* using ^15^N-labeled nitrite and nitrate. **(b)** HRMS analysis of ^15^N incorporation into kinamycin D when no labeled nitrogen-containing salts, 1 mM ^15^N-nitrite, or 1 mM ^15^N-nitrate, were added to the fermentation culture of *S. murayamaensis*. The calculated *m/z* ([M+H]^+^) for kinamycin D at natural abundance is 455.1091, ^15^N-kinamycin D is 456.1055, and ^15^N_2_-kinamycin D is 457.1026. **(c)** 1D ^15^N NMR analysis of kinamycin D isolated from fermentation cultures of *S. murayamaensis* revealed ^15^N-nitrate and ^5^N-nitrite labeled the proximal nitrogen of the diazo group^27^.

### Reconstitution of KinNM activity

To further demonstrate that **3** and acetylhydrazine are also relevant intermediates in the biosynthesis of kinamycin, we first sequenced the genome of *S. murayamaensis* and identified the homologs of FzmNOPQR, which we have labeled as KinLKJNM, respectively. Then we expressed and purified KinN and KinM, which are homologs of FzmQ and FzmR, respectively. We observed the conversion of **3** to **4** by KinN in the presence of acetyl-CoA (**Supplementary Fig. 3ab**). Compound **4** was converted by KinM into acetylhydrazine and fumaric acid as monitored by ^1^H-NMR spectroscopy and LC-HRMS (**Supplementary Fig. 3ac**). We next synthesized doubly-labeled ^15^N_2_-acetylhydrazine and fed this substrate to *S. murayamaensis*. We observed 20% enrichment of both nitrogen atoms of kinamycin D by LC-HRMS ([M+H]^+^ = 457.1026) (**Supplementary Fig. 3d**). These results indicate a role for KinNM and acetylhydrazine in assembling the diazo group of kinamycin D.

### Reconstitution of FzmN and KinL activity

Having confirmed that the nitrogen atoms involved in the hydrazide linkage of fosfazinomycin and the diazo group in kinamycin likely have common origins, we next focused on other enzymes encoded by the conserved five-gene cassette. FzmN is a glutamine synthetase homolog, a class of enzymes that canonically catalyzes the formation of glutamine from glutamic acid and ammonia. We first tested whether FzmN could catalyze this prototypical reaction. FzmN was expressed recombinantly in *E. coli* with an *N*-terminal His_6_-tag. The purified recombinant protein was then incubated with glutamic acid and ammonium chloride in the presence of ATP and MgCl_2_ (**Fig. 4a**). When the reaction was analyzed by ^1^H NMR spectroscopy, a new peak was observed at 2.24 ppm that was absent when FzmN was omitted (**Fig. 4bc**). This new resonance is consistent with the γ-protons on glutamine, and this assignment was confirmed by spiking with a glutamine standard.

**Figure 4.**
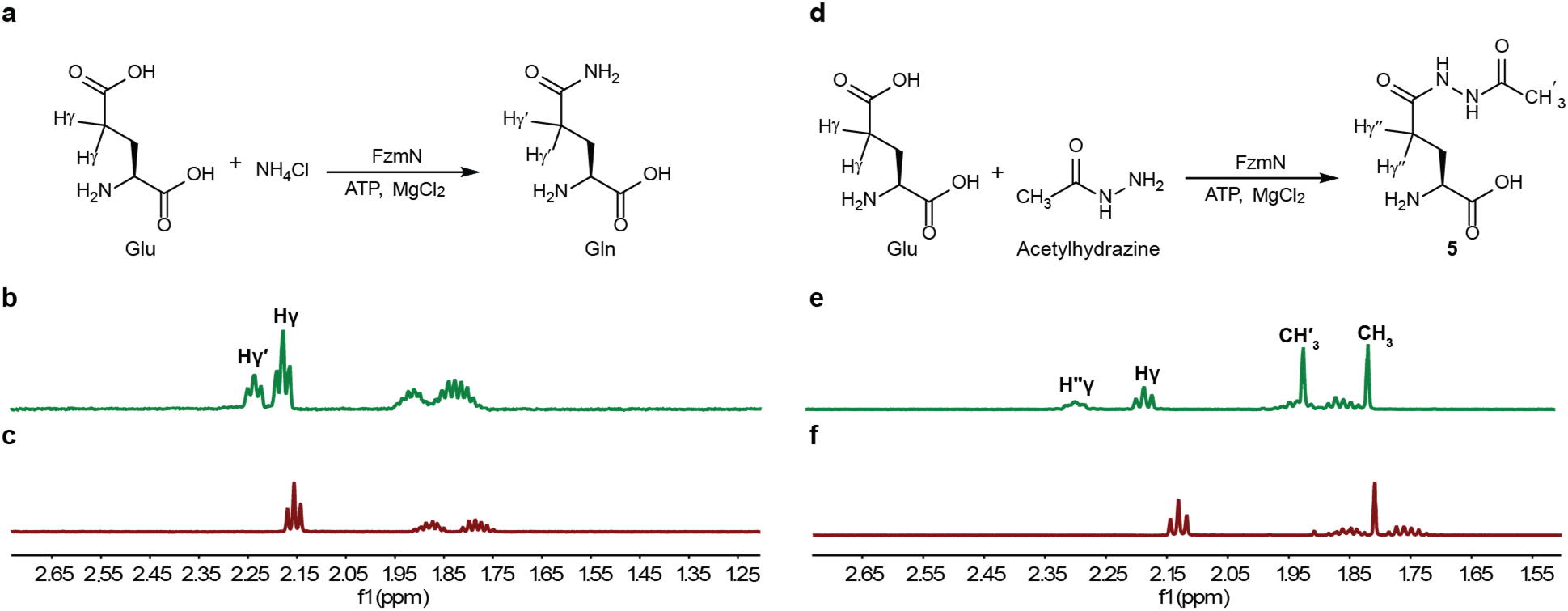
^1^H NMR analysis of FzmN activity. **(a)** The canonical glutamine synthetase reaction catalyzed by FzmN. The γ-protons on the Glu substrate (γ) and Gln product (γ＇) are highlighted. **(b)** ^1^H NMR analysis of the reaction mixture containing FzmN, Glu, NH_4_Cl, ATP, and MgCl_2_. Peaks corresponding to the γ-protons on Glu and Gln are indicated. **(c)** ^1^H NMR analysis of the reaction mixture containing Glu, NH_4_Cl, ATP, and MgCl_2_ without FzmN. **(d)** The putative reaction catalyzed by FzmN during the biosynthesis of fosfazinomycin. The γ-protons on the Glu substrate (γ) and the glutamylacetylhydrazine (**5**) product (γ＂) and the methyl protons on the acetylhydrazine substrate (CH3) and **5** (CH＇3) are indicated. **(e)** ^1^H NMR analysis of the reaction mixture containing FzmN, Glu, acetylhydrazine, ATP, and MgCl_2_. Resonances corresponding to the γ-protons and methyl protons on the substrates and the product are indicated. **(f)** ^1^H NMR analysis of the reaction mixture containing Glu, acetylhydrazine, ATP, and MgCl_2_ without FzmN.

FzmQ and FzmR convert **3** to acetylhydrazine^8^, and the labeling studies described above establish that both of the nitrogen atoms from **3** and acetylhydrazine are incorporated selectively into the phosphonohydrazide linkage of fosfazinomycin. Thus, we reasoned that the next step in the biosynthetic pathway could involve acetylhydrazine as a substrate. Indeed, FzmN catalyzes the condensation of glutamic acid and acetylhydrazine to form glutamylacetylhydrazine (**5**) in the presence of ATP and MgCl_2_ (**Fig. 4d**). When the reaction was analyzed by ^1^H NMR spectroscopy, two new resonances belonging to (**5**) at 2.28 ppm and at 1.92 ppm were observed that were absent when FzmN was withheld from the reaction mixture (**Fig. 4ef**).

Kinetic parameters were determined for FzmN using a coupled enzyme assay with pyruvate kinase and lactate dehydrogenase, measuring NADH consumption by UV-vis spectroscopy. First, the canonical glutamine synthetase reaction with ammonium chloride as a substrate was analyzed, yielding a *k*_cat_ of 0.91±0.02 s^−1^ and a *K*_M_ of 17±1 mM (**Supplementary Fig. 4a**). The FzmN reaction with acetylhydrazine displayed a similar *k*_cat_ (~1.4 s^−1^), but acetylhydrazine was such a good substrate that the *K*_m_ could not be determined and is at least three orders of magnitude lower than the *K*_m_ of ammonia (*K*_m,acetylhydrazine_ < 10 µM) (**Supplementary Fig. 4b**). Together, these results strongly suggest that the condensation of acetylhydrazine and glutamic acid is the role of FzmN in the biosynthesis of fosfazinomycin.

We also tested KinL, the homolog of FzmN encoded in the *kin* biosynthetic gene cluster, for this ligase activity. While we were not able to obtain this enzyme in soluble active form, we confirmed its activity *in vivo* by feeding acetylhydrazine to *E. coli* expressing KinL (**Supplementary Fig. 5a**). Compound **5** was observed in culture supernatants only when KinL was expressed and acetylhydrazine was present in the media (**Supplementary Fig. 5b**). These results demonstrate that KinL possesses analogous reactivity to FzmN.

### Reconstitution of FzmO activity

We next turned our attention towards FzmO, an amidase homolog shared between the biosynthetic pathways of fosfazinomycin and kinamycin. Despite numerous different attempts, we were unable to express recombinant FzmO solubly in *E. coli*. We then endeavored to refold the protein *in vitro*. FzmO, expressed with an *N*-terminal His_6_-tag, was extracted from inclusion bodies with buffers containing guanidine hydrochloride. After purification with nickel affinity chromatography, the protein was diluted into 216 different buffers in 96-well plates^14^. We were able to identify conditions that resulted in solubilized FzmO by measuring the turbidity of the wells with a plate reader. The refolded soluble FzmO was then assessed for enzymatic activity. Glutamyl-acetylhydrazine (**5**), the product of the FzmN-catalyzed reaction, contains an acetyl group that would likely have to be lost prior to funneling the hydrazine synthon into fosfazinomycin biosynthesis. Likewise, in kinamycin biosynthesis, removal of the acetyl group will be necessary before the formation of the final product. Thus, we investigated whether FzmO, being an amidase homolog, might catalyze such a reaction. Refolded FzmO was incubated with **5**, and the products in the reaction mixture were derivatized with fluorenylmethyloxycarbonyl chloride (Fmoc-Cl) to facilitate reversed-phase LC-MS analysis (**Fig. 5a)**. When the reaction mixture was analyzed by LC-MS, a mass consistent with glutamylhydrazine (**6**) was detected that was absent when FzmO was omitted from the reaction (**Fig. 5bc**). To confirm the presence of **6**, a synthetic standard was prepared, and LC-MS analysis revealed that the synthetic material eluted with the same retention time as the product of the FzmO-catalyzed reaction (**Fig. 5d**). The assignment was confirmed by spiking the synthetic standard into the reaction mixture (**Fig. 5e**). The FzmO-catalyzed hydrolysis appears to be selective for the acetylhydrazide linkage over the glutamylhydrazide bond since glutamic acid could not be detected after the reaction (**Fig. 5fg**).

**Figure 5.**
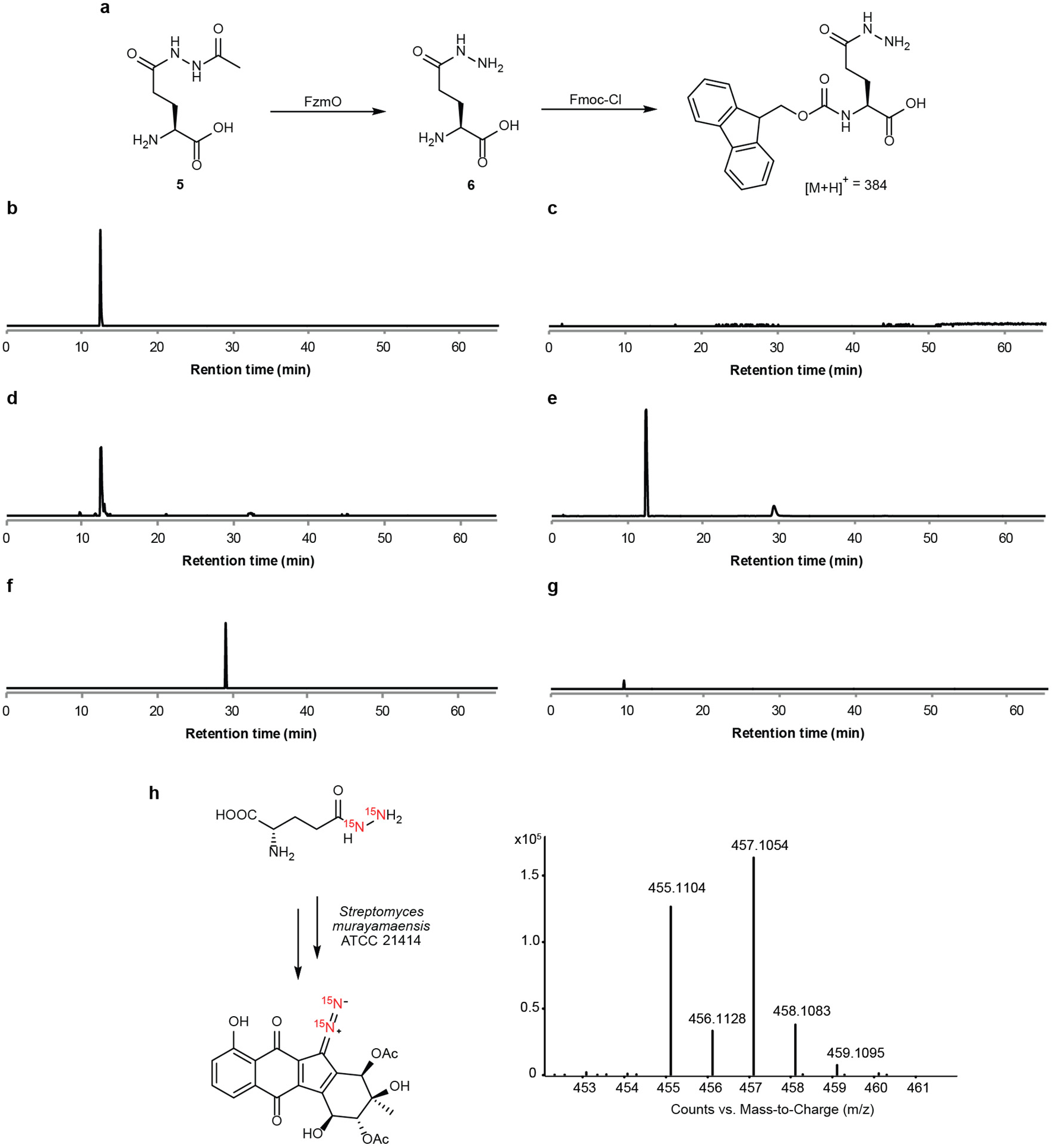
FzmO catalyzes the formation of glutamylhydrazine (**6**) from glutamylacetylhydrazine (**5**), and (**6**) is an intermediate in kinamycin biosynthesis **(a)** Reconstituted deacetylation of **5** to **6** by FzmO. The reaction mixture was derivatized with Fmoc-Cl, and the mass of Fmoc-derivatized **6** ([M+H]^+^ = 384) was monitored by LC-MS. **(b)** Extracted ion chromatogram (EIC) for m/z = 384 of the reaction containing FzmO and **6** in buffer. **(c)** EIC for m/z = 384 of **6** in buffer without FzmO. **(d)** EIC for m/z = 384 of a synthetic standard of **6** in buffer. **(e)** EIC for m/z = 384 of a synthetic standard spiked into the reaction mixture containing FzmO and **6** in panel **b**. **(f)** EIC for m/z = 368 of Fmoc-derivatized Glu in buffer. **(g)** EIC for m/z = 368 of the reaction mixture of FzmO and glutamylacetylhydrazine in buffer. **(h)** HRMS analysis of ^15^N incorporation into kinamycin D from feeding 0.2 mM of ^15^N_2_-**6** to cultures of *S. murayamaensis*. ^15^N_2_-kinamycin D ([M+H]^+^ = 457.1026).

The *kin* gene cluster encodes a homolog of FzmO, KinK. We demonstrated that **6**, the presumed product of KinK-mediated hydrolysis, is an on-pathway intermediate in the biosynthesis of kinamycin D using feeding experiments. We synthesized doubly labeled ^15^N_2_-L-glutamylhydrazide (**6**) and supplemented the fermentation broth of *S. murayamensis* with this compound at a final concentration of 200 µM, as higher concentrations were found to have an adverse effect on growth and production of kinamycin D. We observed 60% enrichment of the diazo group in kinamycin D (**Fig. 5h**). This observation confirms that this functional group derives from a hydrazine-bound glutamyl scaffold.

### Reconstitution of FzmA activity

Having reconstituted the production of **6** by FzmO, we next explored ways in which the hydrazine moiety of **6** could be transferred onto the carboxyl group of arginine since our previous work suggested that argininylhydrazine is an intermediate in the biosynthesis of fosfazinomycin^15^. Such a reaction would minimally require both the breakage and formation of a hydrazide linkage. FzmA, an asparagine synthetase homolog, stood out as a prime candidate to catalyze this reaction. Bifunctional asparagine synthetases contain a glutaminase domain that hydrolyzes the side chain carboxamide of glutamine to liberate ammonia, which travels through a tunnel to a distal synthetase active site where aspartic acid is adenylated and reacted with the ammonia^16^. To assess enzymatic activity, FzmA was expressed recombinantly in *E. coli* bearing a *C*-terminal His_6_-tag since the *N*-terminal cysteine residue is critical for catalysis in canonical asparagine synthetases. After the recombinant protein was incubated with **6,** the products of the reaction mixture were derivatized with Fmoc-Cl. Fmoc-Glu was detected by LC-MS (**Supplementary Fig. 6a**), whereas this product was not observed when FzmA was omitted from the reaction (**Supplementary Fig. 6b**). At present we have not been able to observe condensation of the hydrazine with arginine to form argininylhydrazine, but FzmA clearly has the ability to liberate hydrazine from glutamylhydrazine.

### Free hydrazine is not involved in diazo installation in kinamycin biosynthesis

The fosfazinomycin and kinamycin biosynthetic pathways employ the key intermediate **6** in distinct ways. To construct the diazofluorene scaffold of the kinamycins, the hydrazine moiety of **6** must be hydrolyzed and transferred to a polyketide intermediate. We have previously reported that an electrophilic species generated from the biosynthetic intermediate dehydrorabelomycin by the flavin monooxygenase AlpJ/KinO1 can be attacked non-enzymatically by a variety of amine nucleophiles (**Fig. 6**)^17^. We envisioned that hydrazine liberated from **6** could attack this scaffold to generate a hydrazine adduct, which could be further oxidized to afford the diazofluorene. However, this reaction is unlikely to involve non-enzymatic addition of free hydrazine as we would have observed labeling of both the distal and proximal diazo nitrogen atoms in our ^15^N-nitrite and ^15^N-nitrate feeding experiments (**Fig. 3**).

**Figure 6.**
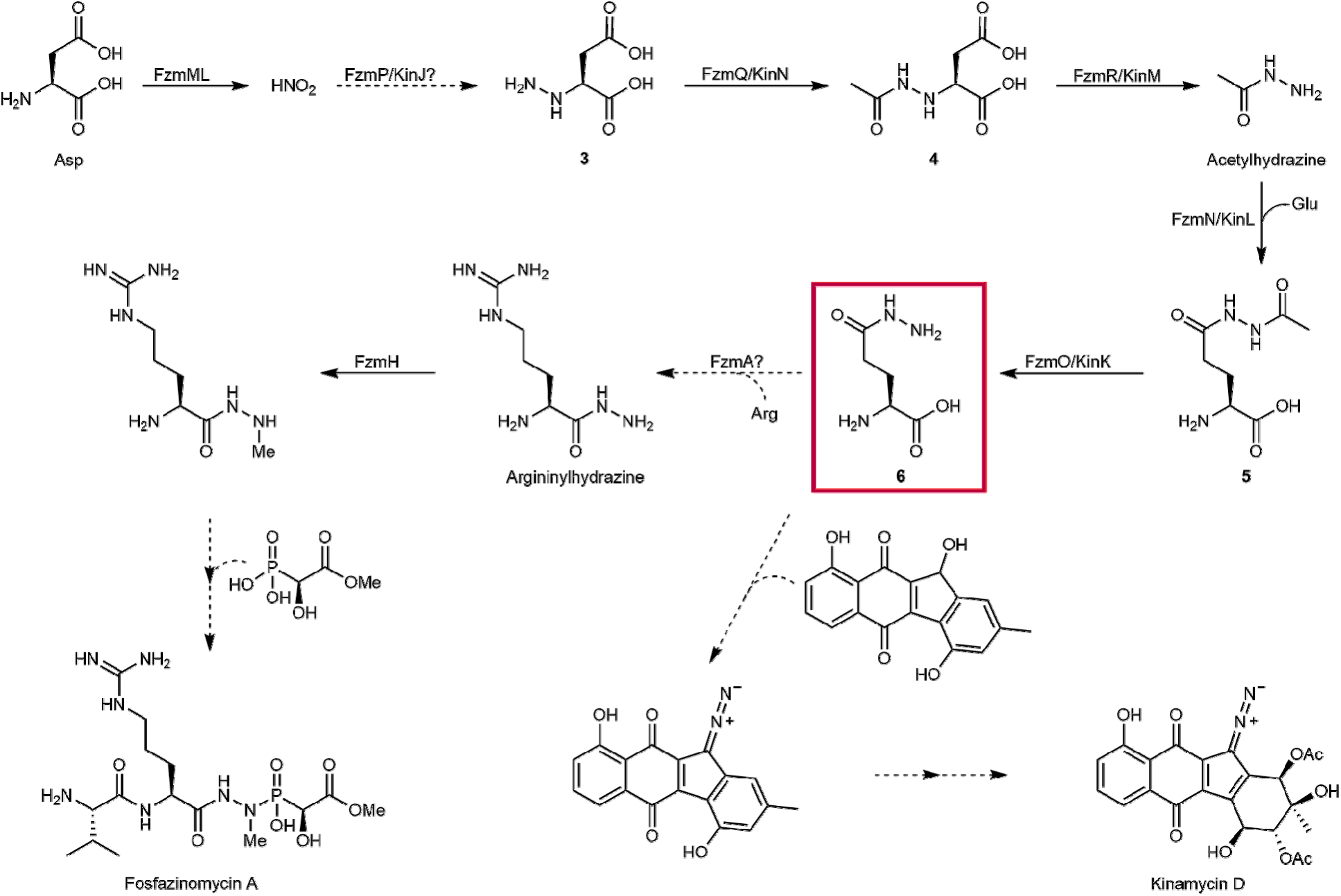
The proposed biosynthetic pathways for fosfazinomycin and kinamycin. Solid arrows indicate steps that have been reconstituted. Dashed lines indicate putative steps. Compound (**6**) acts as the carrier of the hydrazine synthon and is the branch point where the biosyntheses of fosfazinomycin and kinamycin diverge.

To test whether free ^15^N_2_-hydrazine can directly label kinamycin D, we added ^15^N_2_-hydrazine to the fermentation broth of *S. murayamaensis* at a concentration of 0.2 mM, the same concentration at which we fed ^15^N_2_-**6**. At this concentration, we observed 0% labeling (**Supplementary Fig. 7b**). At 1 mM concentration, we observed ~40% labeling of both diazo nitrogen atoms by mass spectrometry (**Supplementary Fig. 7**). We hypothesized that KinL might exhibit promiscuous ligase activity, generating ^15^N_2_-**6** from ^15^N_2_-hydrazine directly in the producing strain. To test this proposal, we fed 1 mM hydrazine to *E. coli* expressing KinL and observed production of **6** only when KinL was expressed and hydrazine was present (**Supplementary Fig. 5c**). Therefore, the labeling we observed from feeding ^15^N_2_-hydrazine likely arises from its incorporation into **6** by KinL.

## Discussion

The *in vivo* feeding experiments presented here demonstrate that one of the nitrogen atoms in the N-N bonds of both fosfazinomycin and kinamycin is derived from nitrous acid, similar to findings in the biosynthesis of cremeomycin. But unlike in the biosynthesis of cremeomycin, for which it has been proposed that the diazo group is formed late in the pathway from a pre-installed aromatic amine, we show that in the biosyntheses of fosfazinomycin and kinamycin, the N-N bond is made early and that the hydrazine functionality is carried through multiple enzymatic steps. Our feeding experiments corroborate the previously reconstituted *in vitro* activities of FzmQ and FzmR, thus firmly establishing that their physiological roles are indeed the acetylation of hydrazinosuccinate and the elimination of acetylhydrazine, respectively (Fig. 6). Further, the reconstitution of the activities of KinN and KinM, the homologs of FzmQ and FzmR in the kinamycin biosynthetic cluster and the results of the feeding studies for kinamycin production illustrate that these steps are conserved in the biosynthetic pathways towards fosfazinomycin and kinamycin. These findings are surprising since previous studies had suggested that the N-N bond in the diazo group of kinamycin was fashioned from an aromatic nitrogen-containing precursor, analogous to the proposal for cremeomycin, but such a pathway is inconsistent with the labeling studies presented here.

To follow the fate of acetylhydrazine, we also reconstituted the activities of FzmN and FzmO, revealing that acetylhydrazine is first condensed onto the side chain carboxyl group of glutamic acid to form glutamylacetylhydrazine (**5**) before deacetylation to yield glutamylhydrazine (**6**). Glutamic acid is present in relatively high concentrations in the cytoplasm of most bacteria, and perhaps for that reason, glutamic acid has previously been demonstrated to be a common carrier molecule to effect many different types of transformations^18^. Most examples come from catabolism, but some have been uncovered in secondary metabolism. For instance, in the biosynthesis of butirosin, an intermediate conjugated to an acyl carrier protein (ACP), γ-aminobutyryl-ACP, is condensed onto the side chain of glutamic acid, and this glutamylated intermediate undergoes two further enzymatic transformations before deglutamylation^18-20^.

In the fosfazinomycin biosynthetic pathway, we previously reported that argininylhydrazine is the substrate for the *N*-methyltransferase FzmH^15^. Thus, the next step in the biosynthesis after formation of **6** likely involves the transfer of hydrazine from the side chain of glutamic acid onto the carboxyl group of arginine. FzmA, an asparagine synthetase homolog, might catalyze such a reaction (**Fig. 6**). Indeed, we have reconstituted the hydrolysis of **6** by FzmA, liberating hydrazine. While at the present, we have yet to reconstitute the enzymatic formation of argininylhydrazine by FzmA, there is precedent for asparagine synthetase homologs catalyzing the transfer of N-X moieties from glutamylated intermediates. A recent report demonstrated that the asparagine synthetase homolog TsnB9 catalyzes the transfer of hydroxylamine from the side chain of glutamic acid to the carboxyl group of an advanced intermediate during the course of trichostatin biosynthesis (**Supplementary Fig. 1e**)^21^.

This work provides further support for a major revision in the proposed biosynthetic pathway for kinamycin and the other diazofluorene polyketides. A previous study had suggested that stealthin C is a biosynthetic intermediate to kinamycin on the basis of isotope labeling studies; however, that study measured the incorporation of deuterated stealthin C, and only low percentages of incorporation were observed (**Supplementary Fig. 1d**)^13^. Furthermore, we have recently reported that stealthin C can be formed nonenzymatically *in vivo* and *in vitro*^17^. Therefore, we posit that stealthin C is not an authentic intermediate in kinamycin biosynthesis, and instead, the N-N bond is formed independently and later added to the polyketide scaffold in a convergent biosynthetic process. Not only do we observe the intact incorporation of the preformed N-N bond into kinamycin, we observe the incorporation of nitrite into the proximal nitrogen in the diazo functionality, which precludes a pathway involving late-stage diazotization. The exact N-N bond-containing intermediate transferred to the polyketide scaffold and the enzymes involved in this process remain to be elucidated.

Collectively, our results show that in the biosyntheses of the structurally highly diverse compounds fosfazinomycin and kinamycin, a hydrazine building block is channeled into the biosynthetic pathways by a common strategy. The N-N bond is first fashioned by conversion of aspartic acid to hydrazinosuccinic acid (**3**) in a process that involves nitrite but that has yet to be fully understood. Then a set of four conserved enzymes transfer the hydrazine moiety onto the side chain of glutamic acid before its final installation onto the mature natural product (**Fig. 6**). This strategy is vastly different than in the biosyntheses of other N-N bond-containing natural products for which the NN bond is formed directly on the scaffold of the final product. Thus, this study brings to light an unexpected new pathway for the incorporation of N-N bonds in natural products.

## Online Methods

### General methods

All reagents and materials for the fosfazinomycin and kinamycin biosynthetic studies were purchased from Sigma-Aldrich or Fisher Scientific unless otherwise noted. DNA oligonucleotides were obtained from Integrated DNA Technologies (IDT) and Sigma-Aldrich. Enzymes used in cloning were procured from New England Biosciences. DNA sequencing was performed by ACGT Inc., Roy J. Carver Biotechnology Center (University of Illinois at Urbana-Champaign), or Eton Bioscience Inc. (Charleston, MA). NMR experiments were carried out on an Agilent 600 MHz with a OneNMR broadband probe or Varian Inova-500 (500 MHz, 125 MHz), and the data was analyzed with MestreNova software. Gene cluster diagrams were constructed with the online Gene Graphics tool^22^. Genome sequencing of *Streptomyces murayamaensis* ATCC 21414 was performed by Era7 Bioinformatics (Granada, Spain) using Illumina MiSeq reads of two 280 bp paired-end libraries^23, 24^. Annotations were carried out using Era7’s tool BG7. Sequence alignments were performed with Geneious. The S. murayamaensis genome has been deposited into NCBI (Accession # TBD)

### Isotope labeling experiments

*Streptomyces* sp. NRRL S-149 was first cultivated on R2A agar and then in ATCC 172 medium as previously described^8^. After 3 d on a roller drum at 30 °C, 1 mL of the culture in ATCC 172 was used to inoculate a second seed culture in 25 mL of modified R2A medium (3.8 mM ^15^NH_4_Cl, [^15^N, 99%, Cambridge Isotope Laboratories], 0.05% soluble potato starch, 2.8 mM glucose, 2.7 mM sodium pyruvate, 0.9 mM potassium phosphate dibasic, and 0.2 mM magnesium sulfate heptahydrate) in a 125 mL flask. After 3 d shaking at 200 rpm at 30 °C, 4 mL of the seed culture was used to inoculate a production culture consisting of 100 mL of modified R2AS (same as modified R2A above, supplemented with 100 µM ^15^N-aspartic acid [15N, 99%, Cambridge Isotope Laboratories], 40 mM sodium succinate, and 0.5% Balch’s vitamins) in a 500 mL baffled flask. The production culture was cultivated for 38 h at 30 °C with shaking at 200 rpm. Then 2 mM of acetylhydrazine, N-hydroxyaspartic acid (**2**), hydrazinosuccinic acid (**3**), or NaNO_2_ were added to the culture. Compounds **2** and **3** were obtained enzymatically from fumaric acid and either hydroxylamine or hydrazine using ammonia-aspartate lyase (AspB) from *Bacillus* sp YM55-1 as previously reported^25^. The production culture was then incubated for an additional 8 h at 30 °C with shaking at 200 rpm. The spent medium was concentrated 10X under reduced pressure and reconstituted in 80% MeOH. Precipitate was removed by centrifugation, and the supernatant was dried under reduced pressure and dissolved in 5 mL of H_2_O. Solid phase extraction with 300 mg of Oasis HLB resin (Waters) was performed; after the sample was loaded onto the resin, the resin was washed with 5% MeOH in H_2_O, and the analyte was eluted with 50% MeOH. LC-HRMS and LC-HR-MS/MS analyses in conjunction with the labeling experiments in *Streptomcyes* sp. NRRL S-149 were performed on a Thermo Q-Exactive Hybrid Quadrupole-Orbitrap Mass Spectrometer coupled to a Dionex Ultimate 3000 series HPLC system. An Xbridge C_18_ column (4.6 x 250 mm, 5 µ, Waters) was used using mobile phase A (H_2_O with 0.1% formic acid) and mobile phase B (acetonitrile with 0.1% formic acid) at a flow rate of 0.5 mL/min. The chromatographic method consisted of a linear gradient of 5% B to 7.5% B in 10 min, a linear gradient from 7.5% B to 95% B in 7 min and a linear gradient from 95% B to 5% B in 11 min. Analyses were performed in positive mode. Data was analyzed and processed with Thermo Xcalibur 3.0.63 software.

*S. murayamaensis* ATCC 21414 was cultivated on mannitol-soy agar for 5 d until sporulation. The spores were inoculated into 50 mL of ISP2 medium for 2 d with shaking at 220 rpm at 30 °C. Then, 1 mL of seed culture was transferred to 100 mL of production medium (glycerol 3%, (NH_4_)_2_SO_4_ 0.1%, K_2_HPO_4_ 3H_2_O 0.1%, CaCO_3_ 0.1%, MgSO_4_ 7H_2_O 0.01%, and FeSO_4_ 7H_2_O 0.01%) containing either ^15^N-sodium nitrite [15N, 98%, Cambridge Isotope Laboratories], ^15^N-calcium nitrate [15N, 98%, Cambridge Isotope Laboratories], ^15^N_2_-hydrazine sulfate [15N, 98%, Cambridge Isotope Laboratories], or ^15^N_2_-acetylhydrazine added to a final concentration of 1 mM. ^15^N_2_-**6** was added to the fermentation culture to a final concentration of 0.2 mM. The fermentation proceeded for 3 d, after which the cultures were centrifuged to remove the cells. The supernatants were extracted twice with ethyl acetate containing 1% acetic acid. The samples were concentrated, redissolved in methanol, and analyzed by LCHRMS using an Agilent 1200 series LC system coupled to an Agilent qTOF 6530 mass spectrometer with a Hypersil Gold aq C_18_ column (3 x 150 mm, 3 µ) using mobile phase A (H_2_O + 0.1% formic acid) and mobile phase B (acetonitrile + 0.1% formic acid) at a flow rate of 0.3 mL/min. The method consisted of a linear gradient from 5% B to 95% B in 15 min, an isocratic hold at 95% B for 5 min, a linear gradient from 95% B to 5% B in 4 min, and a final isocratic hold at 5% B for 5 min. Analyses were performed in positive ion mode, and the extracted ion chromatograms were generated using Agilent Chemstation software with a 5 ppm mass window.

For preparation of ^15^N-labeled kinamycin for NMR analysis, organic extracts from 500 mL of fermentation culture were prepared as described above. Kinamycin D was isolated from the extracts using a ThermoFisher Dionex Ultimate 3000 HPLC system with a Kromasil 100 C_18_ column (10 x 150 mm, 5 µ) at a flow rate of 3 mL/min. The method consisted of a linear gradient from 5% B to 95% B in 30 min, an isocratic hold at 95% B for 10 min, and a linear gradient from 95% to 5% B in 8 min. Kinamycin D eluted at 23 – 25 min. The collected fractions were lyophilized and redissolved in d_6_-DMSO for NMR analysis.

### Cloning

Genomic DNA was isolated from *Streptomyces* sp. XY332, *Streptomyces* sp. WM6372, and *Streptomyces murayamaensis* ATCC 21414 using an Ultraclean Microbial DNA Isolation Kit (Mo Bio) following the manufacturer’s instructions. From purified gDNA, the genes encoding FzmN (*Streptomyces* sp. XY332), FzmO (*Streptomyces* sp. WM6372) and FzmA were amplified using the primers listed in **Supplementary Table 1** and Phusion polymerase. The PCR products were then purified using the QIAquick PCR Purification Kit (Qiagen). pET15b was linearized using NdeI, amplified by PCR with Phusion polymerase (primers listed in **Supplementary Table 1**), and treated with DpnI. *fzmN* and *fzmO* were ligated into the pET15b backbone, and *fzmA* was ligated into pET23b (digested with NdeI and XhoI) using the Gibson Assembly kit from New England Biosciences. *E. coli* DH5α was used for DNA manipulation. The fidelity of the insertion was verified by sequencing.

pET28a and the genes encoding KinK, KinL, and KinM were amplified using the primers listed in **Supplementary Table 1**. The linear vector and the insert were ligated using Gibson Assembly Kit from New England Biosciences. The insertions were verified by sequencing.

The codon-optimized gene encoding AspB from *Bacillus* sp. YM55-1 was ordered from IDT as a gBlock. Following PCR amplification with Q5 polymerase (primers and synthetic gene in **Supplementary Table 1**), *aspB* was ligated into pET15b (digested with NdeI and HindIII) using the HiFi Assembly Mix from New England Biosciences. The fidelity of the insertion was verified by sequencing.

### Protein purification and refolding

For FzmN and FzmA, the protein was expressed and purified in a manner as previously reported^8^. FzmO was expressed using the same methodology as for FzmN. After cell lysis, the insoluble fraction was resuspended in pellet wash buffer (50 mM NaH_2_PO_4_, 300 mM NaCl, 0.10% Triton X-100, pH=7.5) and passed through a 20G syringe needle several times. Again the insoluble fraction was isolated from the supernatant via centrifugation, and the pellet was resuspended in denaturing wash buffer (50 mM NaH_2_PO_4_, 300 mM NaCl, 6 M guanidine HCl, 10 mM imidazole, 10% glycerol, pH=7.5) and incubated at 56 °C for 15 min. After centrifugation, the supernatant was applied to Ni-NTA agarose resin (Qiagen). Nickel affinity purification proceeded similarly as with FzmN, except all buffers also contained 6 M guanidine hydrochloride.

Protein refolding conditions for FzmO were then screened using a platform adapted from another study^26^. Briefly, 100 µL of 216 different buffers was placed into three clear round bottom 96-well plates (Corning) (**Supplementary Table 2**). Each plate also contained 12 positive control wells, each filled with 91 µg bovine serum albumin in 110 µL of denaturing storage buffer (50 mM NaH_2_PO_4_, 300 mM NaCl, 6 M guanidine HCl, 10 mM imidazole, 10% glycerol, pH=7.5); 12 negative control wells containing 91 µg insoluble FzmO in 110 µL of storage buffer (without guanidine hydrochloride) were also included in each plate. Then, 10 µL of a stock solution of 1 mg/mL FzmO in denaturing storage buffer was diluted into each of the 216 buffers in the 96-well plates. The plates were incubated for 12 h at room temperature with gentle agitation. The turbidity of each well was then measured with a Tecan Infinite 200 Pro plate reader, reading absorbance at λ=410 nm and a bandwidth of 9 nm using multiple circle-filled 3×3 reads and 10 flashes per read. Each plate was prepared and analyzed in triplicate on different days. Hits were defined as conditions that gave rise to absorbance readings within three standard deviations of the mean absorbance of the positive control wells.

For AspB, pET15b_*aspB* was used to transform electrocompetent *E. coli* BL21 (DE3) cells. Transformants were cultivated for 12 h at 37 °C in LB with 100 µg/mL ampicillin, and 10 mL of this culture was then used to inoculate a 1 L of culture in LB with 100 µg/mL ampicillin. Cultivation continued at 37 °C, shaking at 220 rpm, until OD_600_ reached between 0.5 and 0.7, and IPTG was added to 100 µM. The culture was grown for an additional 16 h at 18 °C with shaking at 220 rpm. AspB was then purified in the same manner as described for FzmN.

pET28a_*kinN* and pET28a_*kinM* were used to transform chemically competent *E. coli* BL21 (DE3) cells, and the resulting transformants were cultivated overnight in 12 mL of starter cultures at 37 °C in LB with 50 µg/mL of kanamycin. The next day, 10 mL of this starter culture was used to inoculate 1 L of LB medium containing 50 µg/mL of kanamycin, which was allowed to incubate at 37 °C, shaking at 180 rpm, until reaching an OD_600_ of ~0.4-0.6. Then, 200 µM of IPTG was added, and the culture was grown for an additional 16 h at 15 °C. The cells were harvested, resuspended in lysis buffer (20 mM HEPES pH 8.0, 500 mM NaCl) and passed through a cell disruptor. The lysate was clarified via centrifugation, and the resulting supernatant was incubated with Ni-NTA resins for 1 h at 4 °C on a nutating mixer. The subsequent purification steps were analogous to the protocol for purifying FzmN with the exception of the compositions of the wash buffer (50 mM HEPES pH 8.0, 500 mM NaCl, 20 mM imidazole), elution buffer (50 mM HEPES pH 8.0, 500 mM NaCl, 200 mM imidazole), and storage buffer (50 mM HEPES pH 8.0, 50 mM NaCl, 10 mM MgCl_2_, 5% glycerol).

### Enzyme assays

To test whether FzmN could catalyze the canonical glutamine synthetase reaction, 5 mM glutamic acid, 6 mM NH_4_Cl, 50 µM FzmN, 6 mM ATP, and 6 mM MgCl_2_ were incubated in 50 mM NaH_2_PO_4_, pH=7.7 for 5 h at room temperature. The reaction mixture was then treated with Chelex 100 resin for 15 min, and the enzyme was removed by filtration through an Amicon Ultra 30K centrifugal filter (EMD Millipore) before NMR analysis. To test whether FzmN could catalyze the formation of glutamylacetylhydrazine (**5**), 5 mM glutamic acid, 6 mM acetylhydrazine, 34 µM FzmN, 6 mM ATP, and 6 mM MgCl_2_ were incubated in 50 mM NaH_2_PO_4_, pH=7.7 for 8 h at room temperature before Chelex treatment, enzyme removal, and NMR analysis.

To generate Michaelis-Menten curves for FzmN with NH_4_Cl and glutamic acid, a coupled assay system consisting of pyruvate kinase and lactate dehydrogenase was employed and NADH consumption was measured at λ=340 nm with a Cary 4000 spectrometer. The concentration of NH_4_Cl was varied (4 mM to 100 mM) while maintaining a constant concentration of FzmN (1.77 µM). When the concentration of NH_4_Cl was below 10 mM, 100 mM glutamic acid, 36 mM MgCl_2_, 3.75 mM ATP, 1.25 mM phosphoenolpyruvic acid, 200 µM NADH, 50 U/mL pyruvate kinase, and 75 U/mL lactate dehydrogenase were mixed in 50 mM Tris-HCl, pH = 7.7, and the reaction was initiated by adding the substrate, NH_4_Cl. When the concentration of NH_4_Cl used was 10 mM or higher, the same reaction setup was used but with decreased concentrations of pyruvate kinase (30 U/mL) and lactate dehydrogenase (45 U/mL). The attempt to generate kinetic parameters for FzmN towards acetylhydrazine and glutamic acid was executed similarly using the following setup. The concentration of acetylhydrazine was varied (10 µM to 200 µM) while maintaining a constant concentration of FzmN (0.106 µM). 100 mM glutamic acid, 36 mM MgCl_2_, 5 mM ATP, 2 mM phosphoenolpyruvic acid, 200 µM NADH, 40 U/mL pyruvate kinase, and 60 U/mL lactate dehydrogenase were mixed in 50 mM Tris-HCl, pH=7.7, and the reaction was initiated by adding the substrate (acetylhydrazine). Each experiment was performed in triplicate, and the standard deviation was calculated.

In order to test if FzmO is able to deacetylate glutamylacetylhydrazine (**5**), 50 µL was withdrawn from a well in the 96-well plate that was defined as a hit (~10 µM FzmO in 50 mM HEPES, 100 mM KCl, 20% glycerol, 800 mM arginine, pH = 9.0) in the refolding screen. Then, compound **5** was added to a final concentration of 1 mM, and the reaction was allowed to proceed for 11 h at room temperature. The protein was then removed by an Amicon Ultra 10K filter, and the reaction mixture was derivatized with Fmoc-Cl as previously described^8^. A negative control was also run with the same reaction mixture but without any FzmO. After derivatization, the reactions were analyzed by LC-MS using an Agilent 1200 series LC system coupled to an Agilent G1956B single quadrupole mass spectrometer with an Xbridge C_18_ column (4.6 x 250 mm, 5 µ) using mobile phase A (H_2_O) and mobile phase B (acetonitrile) at a flow rate of 1 mL/min. The method consisted of an isocratic hold at 5% B for 1 min, a linear gradient from 5% B to 95% B in 29 min, an isocratic hold at 95% B for 10 min, a linear gradient from 95% B to 5% B in 4 min, and a final isocratic hold at 5% B for 21 min. A positive control was also run wherein 500 µM Fmoc-derivatized glutamylhydrazine was spiked into the reaction mixture. Extracted ion chromatograms were generated using Agilent Chemstation software.

To test the activity of FzmA, 5 mM compound (**6**), 5 mM arginine, 6 mM MgCl_2_, 6 mM ATP, 6 mM thermostable inorganic phosphatase (New England Biosciences), and 21 µM FzmA were incubated in 50 mM NaH_2_PO_4_, pH=7.7 at ambient temperature for 21 h. A negative control was also run using the same conditions as above except that FzmA was withheld from the reaction mixture. After a passage a spin filter to remove enzyme, the reaction mixture was subjected to Fmoc-Cl derivatization as described above before analysis by LC-MS with an Agilent 1200 series LC system coupled to an Agilent G1956B single quadrupole mass spectrometer using a Grace Vydac C_18_ column (4.6 x 250 mm, 5 µ). The method used mobile phase A (H_2_O) and mobile phase B (acetonitrile) at a flow rate of 0.4 mL /min and consisted of an isocratic hold at 20% B for 2 min, a linear gradient from 20% to 100% B in 30 min, an isocratic hold at 100% B for 2 min, a linear gradient from 100% to 20% B in 5 min, and an isocratic hold at 20% B for 5 min. The analysis was performed in negative mode with selected ion monitoring for Fmoc-derivatized glutamic acid ([M-H]^-^ = 368).

To test the activity of KinL, *E. coli* BL21 (DE3) harboring pET28a-*kinL* was grown to OD_600_ ~ 0.5 before adding 0.2 mM IPTG, 1 mM of glutamic acid, and 1 mM of acetylhydrazide or hydrazine. A parallel control with BL21 (DE3) harboring an empty pET28a vector was also prepared. The cells were allowed to grow overnight at 15 °C. The next day, the cultures were centrifuged, and 0.5 mL of the supernatant was added to an equivalent volume of acetone. The samples were centrifuged to remove precipitates and analyzed by LC-HRMS using an Agilent 1200 series LC system coupled to an Agilent qTOF 6530 mass spectrometer. The column used was a Cogent Diamond Hydride column (3 x 150 mm, 4 µ) using mobile phase A (H_2_O + 0.1% formic acid) and mobile phase B (acetonitrile + 0.1% formic acid) at a flow rate of 0.5 mL/min. The method consisted of an isocratic hold at 10% A for 1 min, a linear gradient from 70% A to 10% A in 19 min, an isocratic hold at 70% A for 1 min, a linear gradient from 70% to 10% A in 4 min, and an isocratic hold at 10% A for 4 min.

To test the activity of KinN, 2 mM of hydrazinosuccinic acid, 2 mM of acetyl-CoA, and 20 µM of KinN were mixed in a 200 µL reaction volume with 50 mM potassium phosphate buffer pH 8.0 and 10 mM MgCl_2_. Two negative controls lacking KinN or acetyl-CoA were prepared in parallel. After 1 h, the assay mixtures were frozen and lyophilized to dryness before being resuspended in D_2_O and analyzed by ^1^H-NMR spectroscopy. To test the activity of KinM, KinM was added directly to the NMR sample to a final concentration of 10 µM, and the reaction was monitored by NMR spectroscopy over the course of 1 h.

For LC-HRMS analysis, the same assays were performed in final volumes of 50 µL. The assay mixtures were then analyzed by LC-HRMS in an analogous manner to the procedure for detecting isotopically labeled kinamycin D with the exception that the column used was an Acclaim^TM^ Polar Advantage C_18_ column (ThermoFisher Scientific) (2.1 x 150 mm, 3 µ) and negative ion mode was used. To confirm the consumption of **3** in the activity assay of KinNM, the liquid chromatographic method for detecting **5** was used.

### Chemical synthesis of substrates and standards

The procedures for the chemical synthesis of the various substrates and standards are described in the supplementary Information.

## Acknowledgements

This work was supported by the National Institutes of Health (GM P01 GM077596 to W.A.V. and GM DP2 GM105434 to E.P.B.) a Cottrell Scholar Award (to E.P.B.), and a Camille Dreyfus Teacher-Scholar Award (to E.P.B.), and Harvard University. We thank Dr. Zhong Li of the Metabolomics Laboratory of the Roy J. Carver Biotechnology Center (UIUC) for acquiring HRMS and MS/MS spectra, and Nicholas Lue (Harvard University) for assisting with chemical synthesis. A subset of NMR spectra were acquired on a 600 MHz instrument purchased with support from NIH Grant S10 RR028833.

## Author contributions

Experiments were designed by W.A.V., E.P.B., K.K.A.W, Z.H., T.L.N., and P.W. Experiments were performed by K.K.A.W., T.L.N., P.W., and Z.H. The manuscript was written by K.K.A.W., T.L.N., E.P.B. and W.A.V. The research was supervised by E.P.B. and W.A.V.

